# Van der Waals interactions mediate the enantiomeric substrate preference of the CTP:phosphoglycerate cytidylyltransferase CpgD

**DOI:** 10.64898/2026.05.18.725363

**Authors:** Rideeta Raquib, Tanisha Dhakephalkar, Eric A. Klein, Michael V. Airola

## Abstract

*Caulobacter crescentus* is a gram-negative bacterium that produces the anionic sphingolipid ceramide diphosphoglycerate that can substitute for the lipopolysaccharide component of the outer membrane. *ccna_01210* is a gene in the operon for ceramide diphosphoglycerate synthesis and encodes for the enzyme, CpgD. CpgD is a magnesium-dependent CTP:phosphoglycerate cytidylyltransferase that catalyzes the synthesis of CDP-glycerate, and a pyrophosphate byproduct. CpgD displays substrate specificity for the nucleotide CTP and a preference for 2-phospho-D-glycerate over other phosphoglycerate enantiomers and isomers. Here, we present five high resolution structures for CpgD in various catalytic states that rationalize the specificity and preference of CpgD for its two substrates. This includes structures of apo CpgD, a CpgD-CTP-Mg^2+^ ternary complex, and three CpgD-CDP-glycerate-pyrophosphate-Mg^2+^ product bound complexes. The structures reveal CpgD nucleotide specificity is mediated by favorable hydrogen bonding interactions with the cytosine nucleobase of CTP, while the preference for 2-phospho-D-glycerate occurs due to favorable van der Waals interactions with the 2D enantiomer and unfavorable steric clashes with the 2L enantiomer. A catalytic mechanism involving a pentacoordinate transition state is proposed based on the observed stereochemical inversion of the α-phosphate in the substrate CTP in comparison to the α-phosphate of the product CDP-glycerate. Overall, this provides insights into the catalytic mechanism, nucleotide specificity, and enantiomeric substrate preference of the cytidylyltransferase CpgD that participates in a unique pathway of bacterial sphingolipid synthesis.

## INTRODUCTION

A key feature of gram-negative bacteria is the presence of a lipopolysaccharide (LPS)-rich outer membrane that serves as a barrier against stressors and hydrophobic molecules, making them less susceptible to antibiotics compared to gram-positive bacteria^(1)^. Despite LPS being the first line of defense and a component for survival in most gram-negative bacteria species, LPS-deficient bacterial mutants have been isolated from several species, including *Neisseria meningitidis, Acinetobacter baumanii, Moraxella catarrhalis*, and *Caulobacter crescentus*^(2-5)^. Recent studies with *C. crescentus* demonstrated that it produces the anionic sphingolipid (SL), ceramide diphosphoglycerate (CPG2), with viability partially dependent on CPG2 synthesis in the absence of LPS^(5)^.

SLs are a class of bioactive lipids that contain an amino alcohol sphingosine backbone^(6)^. The eukaryotic sphingolipid pathway has been extensively studied, with ceramide as a central hub^(7)^. SLs have been shown to play a critical role in the regulation of an array of physiological processes in eukaryotes, including inflammation, apoptosis, and cancer metabolism^(7)^. In prokaryotes, some bacteria can utilize host SLs, whereas various bacterial species can produce SLs of their own^(8)^. Several recent studies have characterized prokaryotic SLs that vary in lipid-headgroups, acyl chain lengths, branching, and saturation^(8, 9)^. These SLs play a role in a multitude of bacterial physiological processes, including the preservation of outer membrane integrity, and host-microbe interactions^(8-11)^.

The reliance of *C. crescentus* on anionic SLs for viability in the absence of LPS led to the discovery of a unique bacterial ceramide polyphosphoglycerate pathway^(5)^. Via genetic and biochemical studies, enzymes in the synthesis of ceramide have been established^(12, 13)^. In addition, several enzymes that participate in the synthesis of the polyphosphate glycerate headgroup of CPG2 have been identified or characterized^(5, 14)^. This includes the ceramide kinase, CpgB, encoded by *ccna_01218*, which phosphorylates ceramide to ceramide-1-phosphate (C1P)^(15)^, and CpgC (encoded by *ccna_01219*) and CpgA (encoded by *ccna_01217*) that are respectively involved in the conversion of C1P to CPG, or CPG to CPG2^(5)^ (Fig. 1A).

**Figure 1.**
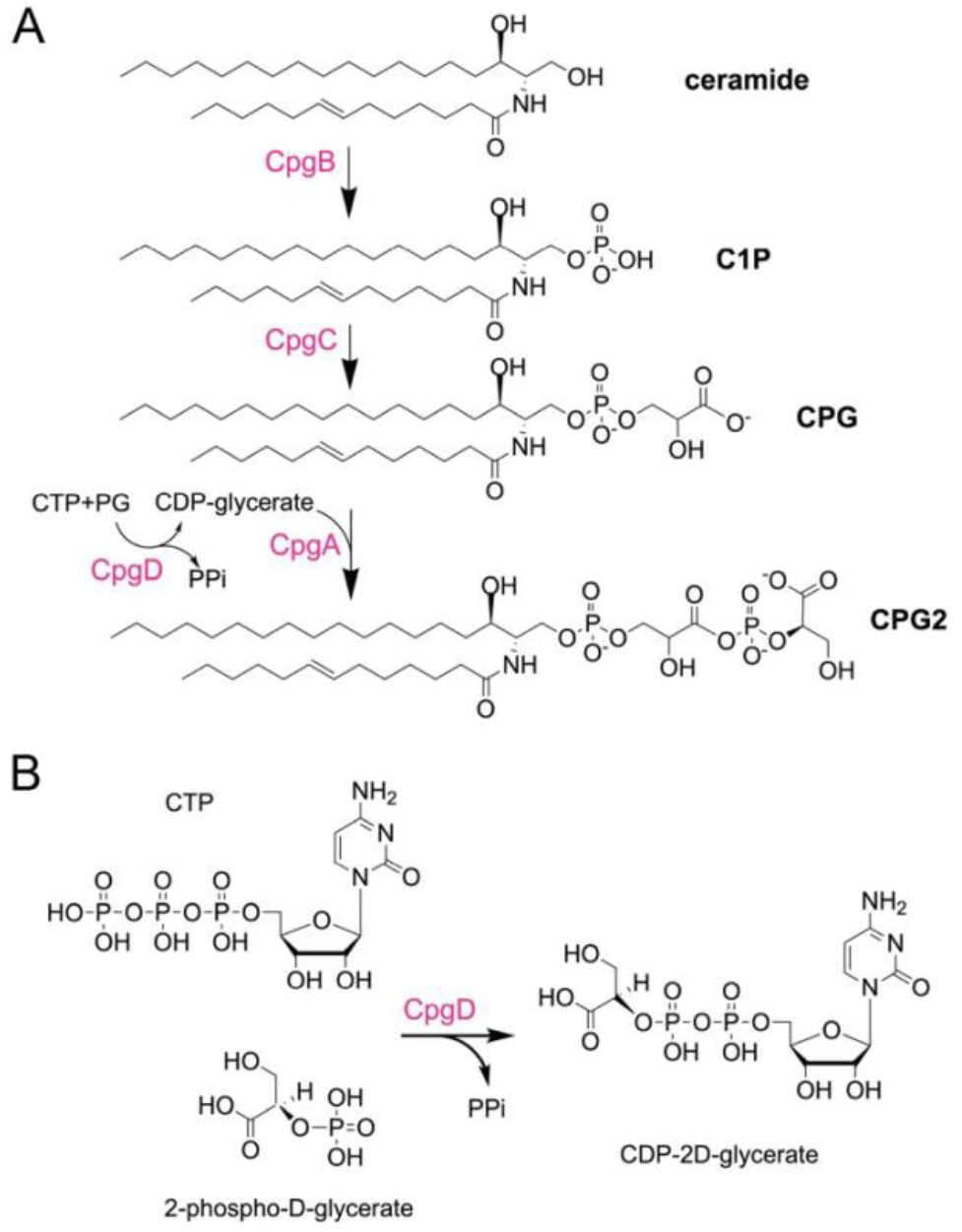
Biosynthesis pathway of CPG2. **A**. Based on previous genetic^5^ and biochemical^11^ studies, a proposed pathway for ceramide diphosphoglycerate (CPG2) synthesis is shown. CpgB is a kinase that uses ATP to convert ceramide to ceramide-1-phosphate (C1P). CpgC, via an unidentified mechanism, is involved in the conversion C1P to ceramide phosphoglycerate (CPG). CpgD converts CTP and phosphoglycerate (PG) to pyrophosphate (PPi) and CDP-glycerate, which is used as a substrate by CpgA to convert CPG to CPG2. **B**. Reaction diagram for the catalytic conversion of CTP and 2-phospho-D-glycerate to CDP-glycerate and pyrophosphate (PPi) by the cytidylyltransferase CpgD.

CpgD (encoded by *ccna_01210*) was recently characterized as a cytidylyltransferase that catalyzes the conversion of CTP and phosphoglycerate (PG) to CDP-glycerate and pyrophosphate^(16)^ (Fig. 1B), with the CDP-glycerate product of CpgD proposed to act as a substrate for CpgA in the subsequent steps of CPG2 synthesis (Fig. 1A). Biochemically CpgD has a strict specificity for CTP over other nucleotides^(16)^ and prefers 2-phosphoglycerate over 3-phosphoglycerate as a substrate. This substrate specificity extends beyond the position of the phosphate group of phosphoglycerate with CpgD displaying a higher affinity and catalytic efficiency for the 2-phospho-D-glycerate enantiomer over 2-phospho-L-glycerate^(16)^.

Here, we present five high resolution structures of CpgD in its apo form, bound to its substrates, Mg^2+^ ions, and products. These structures provide a rationale for the specificity of CpgD for CTP, its preference towards the 2-phospho-D-glycerate enantiomer and suggest a catalytic mechanism involving a penta-coordinate transition state. Overall, this provides new insights into the structural basis of substrate specificity, preference and catalysis by the bacterial cytidylyltransferase, CpgD.

## RESULTS

### Structures of CpgD reveal conformational changes induced by nucleotide binding

To gain insights into CpgD’s catalytic mechanism, substrate preferences, and conformational changes during catalysis, we determined high resolution structures of CpgD in different catalytic states, starting with the apo and CTP bound states. CpgD was overexpressed in *Escherichia coli* and purified using Ni-NTA affinity and size-exclusion chromatography. An apo CpgD structure was determined at pH 4. A CTP and magnesium (Mg^2+^) bound structure was determined at neutral pH, where the enzyme exhibited optimal activity *in vitro*^12^. We were unable to determine an apo structure at neutral pH or a CTP bound structure at acidic pH. The apo and CTP bound structures were refined to 1.8 Å and 1.1 Å resolutions, respectively (Table 1).

**Table 1.** Data Collection and Refinement Statistics.

| **Data Collection and Refinement Statistics** |  |  |  |  |  |
| --- | --- | --- | --- | --- | --- |
|  | apo | CTP Mg | CDP-2D-glycerate | CDP-2L-glycerate | CDP-3D-glycerate |
| PDB ID | 10WG | 10WH | 10WI | 10WJ | 10WK |
| Wavelength, Å | 0.92019 | 0.92019 | 0.9199 | 0.97934 | 0.9793 |
| Resolution range, Å | 34.43 - 1.80<br>(1.84 - 1.80) | 32.48 - 1.10<br>(1.11 - 1.10) | 34.46 - 1.44<br>(1.46 - 1.44) | 41.07 - 1.46<br>(1.49 - 1.46) | 33.5 - 1.51<br>(1.53 - 1.51) |
| Space group | P 21 21 21 | C 1 2 1 | P 21 21 21 | C 2 2 21 | P 21 21 21 |
| Unit cell | 38.8, 77.2, 153.8<br>90 90 90 | 63.1 58.0 77.8<br>90 111.3 90 | 52.9 70.7 153.0<br>90 90 90 | 52.8 72.2 153.4<br>90 90 90 | 54.3 67.3 154.5<br>90 90 90 |
| Total reflections | 368083 (17225) | 713429 (27203) | 1419844 (68985) | 689533 (34790) | 1216824 (61189) |
| Unique reflections | 43786 (2692) | 105101 (3263) | 103790 (3378) | 50903 (2820) | 89542 (2918) |
| Multiplicity | 8.4 (8.0) | 6.8 (5.4) | 13.7 (13.4) | 13.5 (13.7) | 13.6 (13.9) |
| Completeness (%) | 100.0 (100.0) | 99.4 (96.0) | 100.0 (100.0) | 100.0 (100.0) | 99.7 (99.2) |
| Mean I/sigma(I) | 8.6 (1.3) | 12.3 (0.8) | 9.4 (1.0) | 10.0 (1.3) | 9.0 (1.7) |
| Wilson B-factor | 21.2 | 14.04 | 13.37 | 14.94 | 10.07 |
| R-merge | 0.179 (1.861) | 0.056 (1.679) | 0.216 (3.062) | 0.172 (2.434) | 0.337 (3.554) |
| R-meas | 0.191 (1.993) | 0.061 (1.858) | 0.224 (3.182) | 0.179 (2.527) | 0.350 (3.689) |
| R-pim | 0.066 (0.706) | 0.023 (0.786) | 0.060 (0.861) | 0.048 (0.672) | 0.094 (0.981) |
| CC1/2 | 0.998 (0.437) | 0.999 (0.390) | 0.999 (0.324) | 0.998 (0.385) | 0.993 (0.345) |
| CC* | 1 (0.78) | 1 (0.752) | 1 (0.726) | 1 (0.812) | 1 (0.87) |
| Reflections used in refinement | 43761 (2692) | 104945 (3260) | 103756 (3378) | 50881 (2820) | 89256 (2921) |
| Reflections used for R-free | 2177 (141) | 5243 (125) | 5195 (167) | 2552 (143) | 4491 (142) |
| R-work | 0.1863 (0.3193) | 0.1748 (0.4617) | 0.1596 (0.3300) | 0.1723 (0.3750) | 0.1603 (0.2770) |
| R-free | 0.2171 (0.3673) | 0.1903 (0.4289) | 0.1883 (0.3537) | 0.1979 (0.3950) | 0.1866 (0.2970) |
| Number of non-hydrogen atoms | 4348 | 2230 | 4538 | 2312 | 4702 |
| macromolecules | 3977 | 1971 | 3939 | 1990 | 3983 |
| ligands | 13 | 38 | 107 | 50 | 88 |
| solvent | 358 | 221 | 492 | 272 | 631 |
| Protein residues | 510 | 247 | 500 | 250 | 500 |
| RMS(bonds) | 0.011 | 0.011 | 0.124 | 0.016 | 0.007 |
| RMS(angles) | 1.02 | 1.2 | 1.61 | 1.44 | 0.87 |
| Ramachandran favored (%) | 98.21 | 98.37 | 98.19 | 97.98 | 98.19 |
| Ramachandran allowed (%) | 1.59 | 1.63 | 1.61 | 1.61 | 1.61 |
| Ramachandran outliers (%) | 0.2 | 0 | 0.2 | 0.4 | 0.2 |
| Rotamer outliers (%) | 0.24 | 0.47 | 0.47 | 0.47 | 0.47 |
| Clashscore | 1.63 | 1 | 2.11 | 1.97 | 1.73 |
| Average B-factor | 29.1 | 25.06 | 20.46 | 24.45 | 16.82 |
| macromolecules | 28.47 | 23.77 | 18.47 | 22.53 | 14.53 |
| ligands | 25.7 | 20.84 | 26.83 | 21.46 | 13.51 |
| CDP-glycerate | - | - | 10.83 | 18.56 | 12.24 |
| CDP moiety | - | - | 10.49 | 14.98 | 7.89 |
| glycerate moiety | - | - | 12.12 | 33.52 | 30.36 |
| solvent | 36.23 | 37.33 | 36.07 | 39.05 | 31.73 |

Statistics for the highest resolution shell are shown in parentheses.

The structures revealed CpgD to adopt a similar structure to other nucleotidyltransferases^(17-20)^ with an α/β/α fold composed of anti-parallel β-sheets sandwiched between a series α-helices (Fig. 2A). A surface model of the apo CpgD structure had a prominent central cavity, where the active site of the enzyme resides (Fig. 2B). The active site consists of extensive loops that flank the central cavity where the substrates CTP and phosphoglycerate would bind.

**Figure 2.**
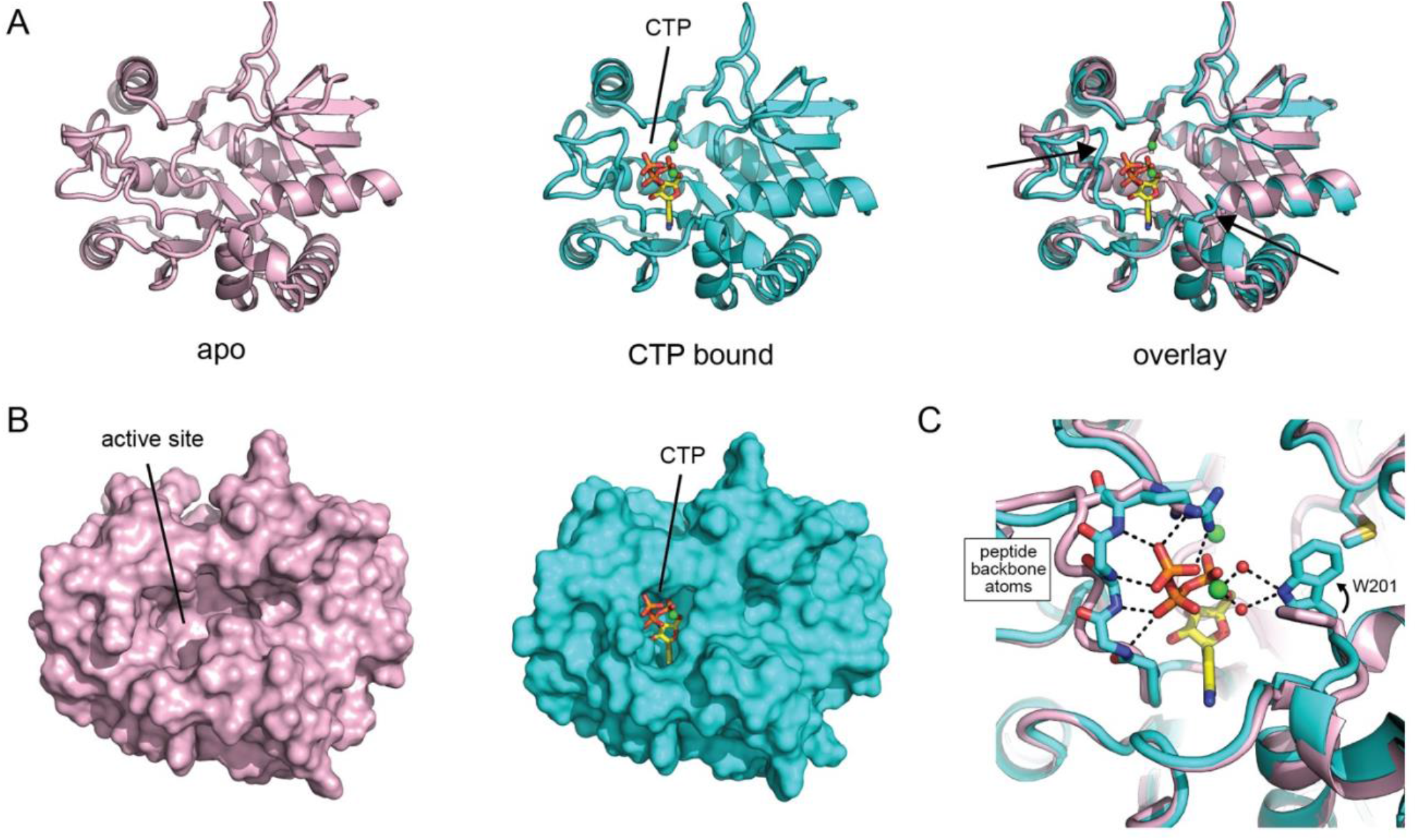
Conformational changes induced by CTP binding. **A**. Cartoon views of the apo (pink) and CTP bound (cyan) structures of CpgD that are superimposed (right). CTP binding induces movement of two loop regions (black arrows) in the active site whose residues are involved in CTP coordination. The carbon (yellow), nitrogen (blue), oxygen (red), and phosphorous (orange) atoms of CTP are colored accordingly. Mg^2+^ ions, green spheres. **B**. Surface views of apo (left, pink) and CTP bound (right, cyan) highlighting the conformational changes within the active site that are induced by CTP binding. **C**. Detailed view of the interactions between CTP and CpgD involved in the active site conformational changes. The backbone nitrogen atoms of “loop 1” move inward to hydrogen bond with the phosphoryl-oxygen atoms of CTP. Trp201 in “loop 2” changes its rotamer position to hydrogen bond with two water molecules that coordinate the central Mg^2+^ ion. Water molecules, red spheres. Bonding interactions, black dotted lines.

In the CTP bound structure, CTP and two Mg^2+^ ions were bound in the active site and were coordinated by a set of conserved residues in other nucleotidyltransferases^(17-19)^. The active site cavity adopted a more closed conformation compared to the apo structure (Fig. 2B). These conformational changes were driven by the movement of two loops whose residues interacted with CTP (Fig. 2A). One of the loops contains Arg16, a highly conserved residue, known to play a role in catalysis^(17-20)^ in the cytidylyltransferase family, which interacted with CTP. The movement of this same loop also resulted in hydrogen bond interactions between the backbone nitrogen atoms of the loop and the phosphoryl-oxygen atoms of CTP (Fig. 2C). Movement of the second loop involved residue Trp201, which switched its rotamer position upon CTP binding to hydrogen bond with two water molecules that coordinated one of the Mg^2+^ ions. Overall, CTP binding resulted in relatively minor conformational changes and was limited to two specific regions that were involved in interactions with either CTP or the Mg^2+^ ions.

### Favorable hydrogen bonding interactions rationalize the nucleotide specificity of CpgD

Electron densities for the CTP molecule (Fig. 3A) and surrounding residues were well resolved in our structure. CTP was coordinated by a series of interactions with the enzyme and two Mg^2+^ ions (Fig. 3B). These interactions involved the conserved Arg16 residue that coordinated the oxygen atoms of the α- and γ-phosphates of CTP (Fig. 3B). Two Mg^2+^ ions were hexa-coordinated through interactions with the phosphoryl-oxygen atoms of CTP, negatively charged acidic residues of CpgD, and water molecules (Fig. 3B). Additional interactions were observed with the 2’ and 3’ hydroxyl groups on the ribose moiety of CTP forming H-bonds with the backbone nitrogen atoms of glycine residues, Gly12 and Gly111, and the O atom within the ribose ring acting as a H-bond acceptor to interact with the sidechain of Asn90 (Fig. 3B).

**Figure 3.**
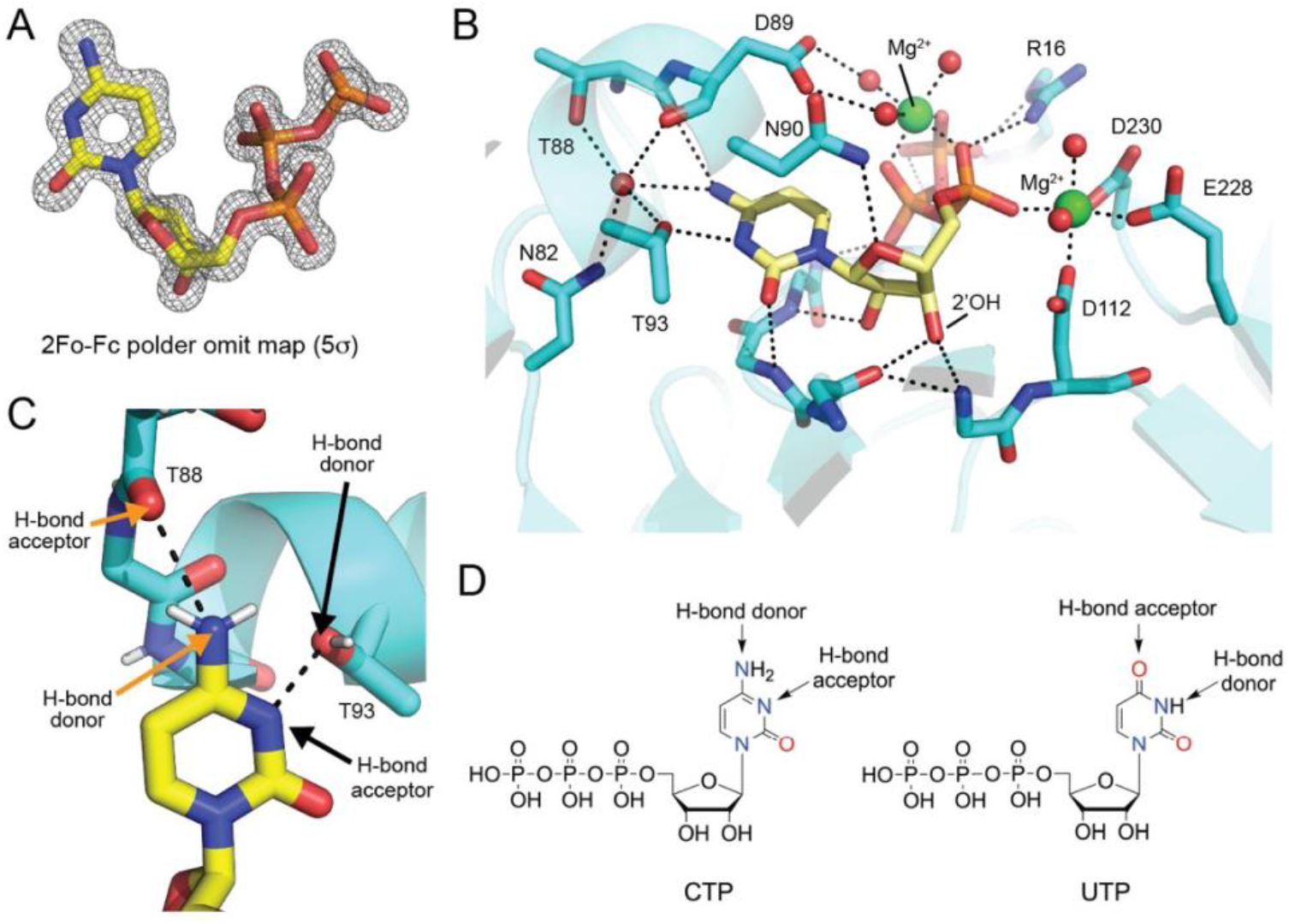
Favorable hydrogen bonding interactions rationalize CTP nucleotide specificity. **A**. 2F_o_-F_c_ electron density for CTP from a polder omit map contoured at 5σ. The CTP molecule was omitted when generating the polder map. **B**. Detailed view of the interactions between CpgD and CTP. CpgD forms extensive interactions with the cytosine ring, ribose, and triphosphate moeities of CTP. Two Mg^2+^ ions aid in the coordination of the CTP phosphates. Hydrogen and ionic bonds are shown by dashes lines. Mg^2+^ ions, green spheres. Water molecules, red spheres. The carbon, oxygen, and nitrogen atoms of CpgD are colored cyan, red, and blue. **C**. The specificity of CpgD for CTP is driven by favorable H-bonding interactions between the sidechain of Thr93 and the carbonyl-oxygen atom of Thr88 with atoms in the cytosine nitrogenous base of CTP. The H-bond donor and acceptor are highlighted for each hydrogen bond interaction (black dotted line). **D**. Comparison of the chemical structures of CTP and UTP that highlights the atoms in the cytosine and uridine pyrimidine rings with differing hydrogen bond acceptor and donor capabilities.

Importantly, the structure allowed us to rationalize the specificity of CpgG for CTP over other nucleotides^(16)^. Examination of the interactions with the cytosine nucleobase of CTP identified two key interactions involving Thr88 and Thr93 (Fig. 3C). Specifically, the amine nitrogen attached to the 4 position of cytosine acted as an H-bond donor to form a hydrogen bond with the backbone carbonyl oxygen of Thr88 (Fig. 3C). Similarly, the nitrogen in the 3 position of the cytosine ring, which is unprotonated, acted as a H-bond acceptor to form a hydrogen bond with the sidechain hydroxyl group of Thr93 (Fig. 3C). These favorable H-bond interactions are only possible with CTP and would not occur with UTP. This derives from the fact that the uridine nucleobase of UTP contains atoms that, in the analogous positions of cytosine, are capable of hydrogen bonding, but with opposing capabilities to act as a hydrogen bond donor or acceptor (Fig. 3D). Overall, the favorable hydrogen bonding interactions provide a clear rationale for CpgD’s nucleotide specificity for CTP^(16)^.

### Structural snapshots of CpgD-mediated catalysis suggest a pentacoordinate transition state

To gain further insight into the catalytic mechanism and substrate specificity of CpgD, we next co-crystallized CpgD with CTP, magnesium, and the preferred enantiomeric substrate, 2-phospho-D-glycerate. This structure was refined to 1.44 Å resolution (Table 1). The structure revealed CpgD bound to the two products, CDP-2D-glycerate and pyrophosphate, and four magnesium ions in the active site (Fig. 4A,B) indicating catalysis had occurred. Overall, there were minimal conformational changes of the active site when comparing the products and CTP bound states (Fig. 4A,C). This suggested that once bound to CTP, the enzyme is poised to bind 2-phospho-D-glycerate and initiate catalysis.

**Figure 4.**
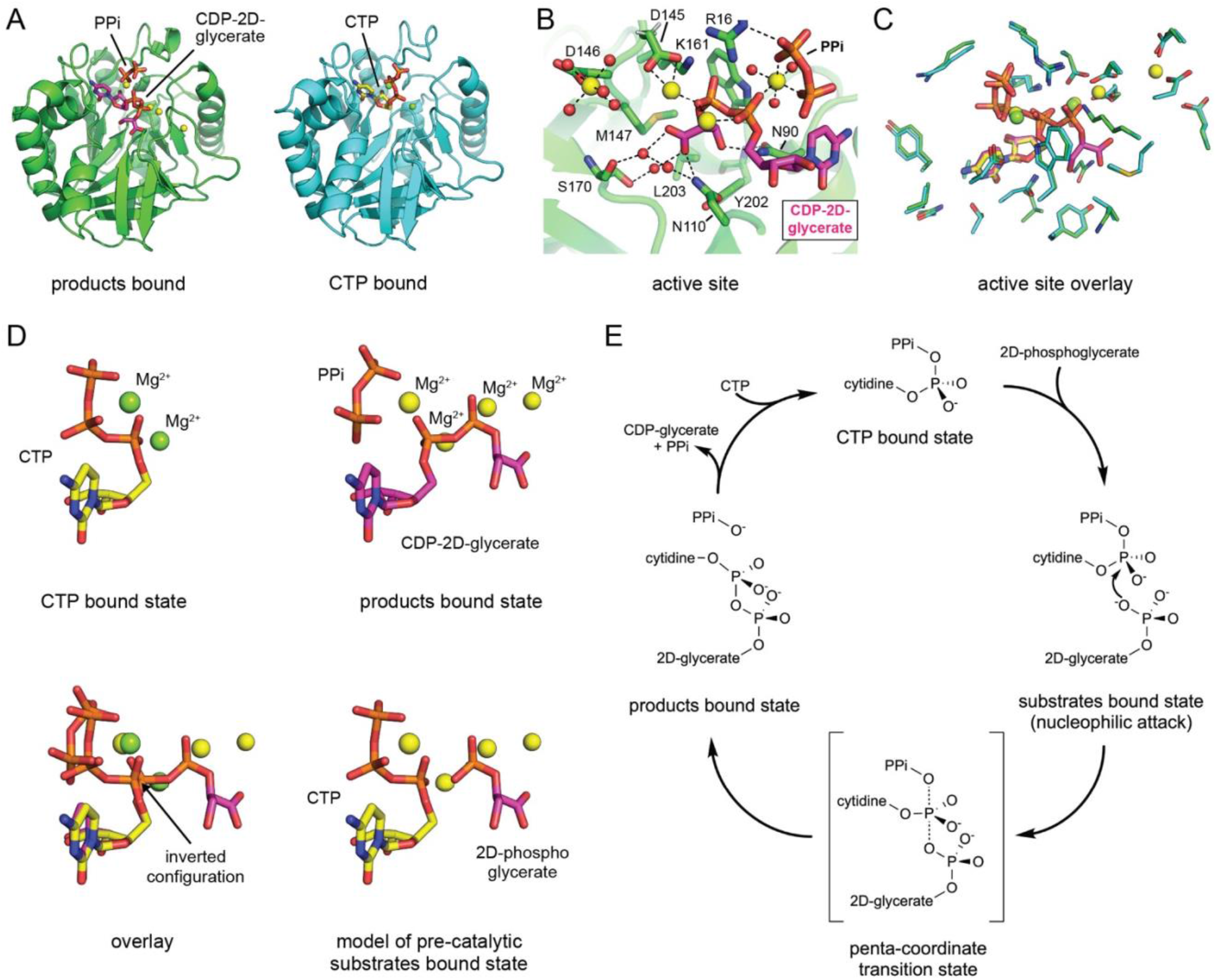
CpgD mediated catalysis proceeds through a penta-coordinate transition state. **A**. Cartoon view of the products bound (green) and CTP bound (cyan) structures of CpgD. The carbon atoms of CDP-2D-glycerate and CTP are colored magenta and yellow, respectively. The nitrogen (blue), oxygen (red), and phosphorous (orange) atoms of CDP-2D-glycerate, CTP, and pyrophosphate (PPi) are colored accordingly. Mg^2+^ ions, yellow or green spheres. **B**. Detailed view of the interactions between CpgD and CDP-2D-glycerate. CpgD forms extensive interactions with the cytosine ring, ribose, diphosphate, and glycerate moieties of CDP-2D-glycerate. Three of the four Mg^2+^ ions are involved in coordinating the phosphate groups of either CDP-2D-glycerate and/or pyrophosphate. Hydrogen and ionic bonds are shown by dashes lines. Mg^2+^ ions, yellow spheres. Water molecules, red spheres. The carbon, oxygen, and nitrogen atoms of CpgD are colored green, red, and blue. **C**. Comparison of the active site residue sidechains in the CTP and products bound structures. All sidechains are in identical positions indicating there are no major conformational changes during catalysis. **D**. Stick representations of CTP (top left), CDP-2D-glycerate and pyrophosphate (PPi) (top right) that are overlayed (bottom left). A stereochemical inversion around the α-phosphate suggests a penta-coordinate transition state during catalysis. A model of the substrates bound state (bottom right) generated by superposition. **E**. Proposed catalytic mechanism for the generation of CDP-2D-glycerate by CpgD. The penta-coordinate transition state is generated by an SN2 nucleophilic attack of the alpha-phosphate group of CTP by an oxygen atom of 2D-phosphoglycerate.

A series of interactions were observed between the CDP-2D-glycerate product and CpgD enzyme that provided insight into how CpgD interacts with the product (Fig. 4B), which are presumably shared with the 2-phospho-D-glycerate substrate. The cytosine and ribose moieties of CDP-2D-glycerate made similar interactions as observed with CTP. The glycerate moiety of CDP-2D-glycerate formed direct hydrogen bonding interactions with Asn90 and Asn110 and was involved in a hydrogen bond network with Ser170 via two bridging water molecules (Fig. 4B).

The product bound structure contained two additional magnesium ions (Fig. 4A,B), for a total of four magnesium ions bound in the active site, in comparison to the two magnesium ions present in the CTP bound structure (Fig. 4A,C). The positions of the two magnesium ions that were also present in the CTP bound state, remained in similar positions and coordinated either the α- and β-phosphates of CDP-2D-glycerate, or bridged the pyrophosphate product and α-phosphate of CDP-2D-glycerate (Fig. 4B). The third newly observed magnesium ion coordinated the β-phosphate of the CDP-2D-glycerate (Fig. 4B), which suggests a potential role in catalysis. The fourth magnesium ion was bound in the active site pocket (Fig. 4B) but was not involved in coordinating either of the CDP-2D-glycerate or pyrophosphate products, making its role unclear.

Comparing the products versus the CTP bound structures suggested the catalytic mechanism of CpgD to involve a penta-coordinate transition state (Fig. 4D,E). In the product bound state, the bond between the α- and β-phosphates of CTP is broken, resulting in the generation of pyrophosphate, which occupies a similar position to the β- and γ-phosphates of CTP prior to the reaction (Fig. 4D). The orientation of the product, CDP-2D-glycerate, provides insights on how the substrate, 2-phospho-D-glycerate, would be positioned pre-catalysis, with a phosphoryl oxygen atom of 2-phospho-D-glycerate oriented for nucleophilic attack on the α-phosphate of CTP (Fig. 4D). Notably, the α-phosphate undergoes a stereochemical inversion if we compare the α-phosphate of the substrate CTP with the α-phosphate of the product CDP-2D-glycerate (Fig. 4D). This strongly suggests that the α-phosphate adopts a penta-coordinate transition state during CpgD mediated catalysis (Fig. 4E). Taken together, we propose the following catalytic mechanism: 1) CTP binds, 2) 2-phospho-D-glycerate binds and 3) the oxygen atom of the phosphate in 2-phospho-D-glycerate carries out an S_N_2 nucleophilic attack on the α-phosphate of CTP, 4) this results in a penta-coordinate transition state, which upon break down, 5) forms the two products CDP-2D-glycerate and pyrophosphate (Fig. 4E).

### Structural basis for the 2-phospho-D-glycerate enantiomeric preference of CpgD

*In vitro* CpgD displays a preference for 2-phospho-D-glycerate as a substrate, when compared to 2-phospho-L-glycerate and the related 3-phospho-D-glycerate^(16)^ with K_M_,_app_ values of 0.8, 3.1, and 10.9 mM, respectively (Fig. 5A). To understand the structural basis for this enantiomeric preference, we determined structures of the product bound states of CpgD after co-crystallization with CTP, magnesium, and either 2-phospho-L-glycerate or 3-phospho-D-glycerate (Table 1). This resulted in CpgD structures bound to the products, pyrophosphate and either CDP-2L-glycerate or CDP-3D-glycerate (Fig. 5B).

**Figure 5.**
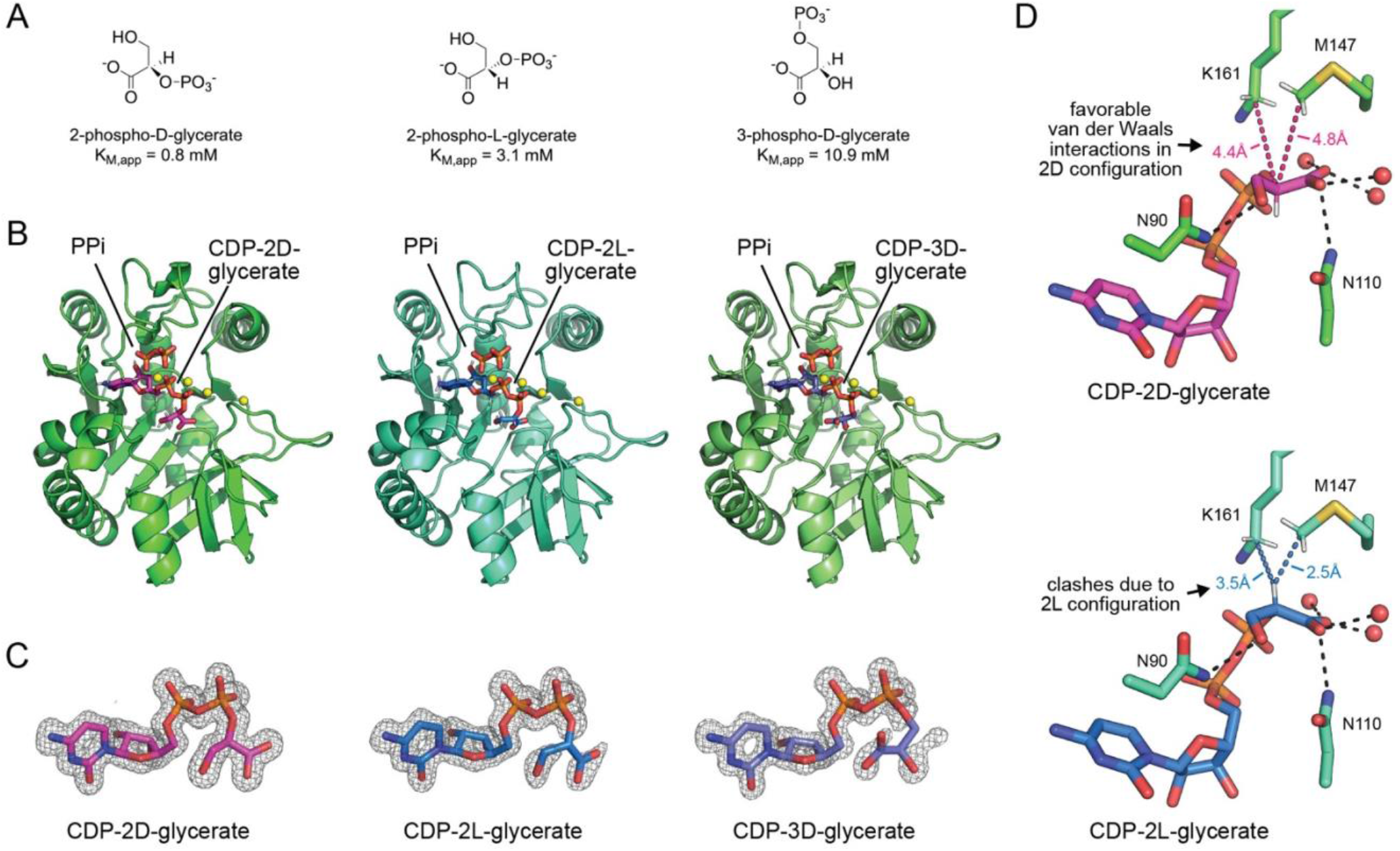
Structural basis for the 2-phospho-D-glycerate enantiomeric preference of CpgD. **A**. Chemical structures and apparent Michaelis-Menten constants (K_M,app_) of the three phosphoglycerate isomers and enantiomers that can be used as substrates by CpgD. **B**. Cartoon views of CpgD structures bound to the products pyrophosphate (PPi) and different isomers of CDP-glycerate. **C**. 2Fo-Fc electron density for the distinct CDP-glycerate products generated from a polder omit map contoured at 5σ. The density varies in the glycerate portion of the CDP-glycerate molecule based on the affinity of the phosphoglycerate substrate. **D**. The specificity towards the 2D enantiomer of phosphoglycerate is mediated by favorable van der Waals interactions with the sidechains of Met147 and Lys161, which in the 2L enantiomer clash with the hydrogen atom that points upward toward these sidechains.

A comparison of the three CpgD product bound structures revealed near identical structures, regardless of the phosphoglycerate isomer used as a substrate, with respect to the position of pyrophosphate and CDP-glycerate products, and the number of active site magnesium ions (Fig. 5B). However, in contrast to CDP-2D-glycerate, where the electron density was clearly defined for the entire molecule (Fig. 5C), the electron density for the glycerate moiety was not clearly defined in the CDP-2L-glycerate or CDP-3D-glycerate structures (Fig. 5C). This was reflected by a contrast between the B-factors of the glycerate moiety, relative to the CDP portion, within the different CDP-glycerate isomers. For CDP-2D-glycerate, the average B-factors for atoms in the glycerate moiety were similar to the CDP portion (Table 1), which indicates the glycerate moiety was well ordered and in a single conformation. In contrast, for CDP-2L-glycerate and CDP-3D-glycerate the B-factors for glycerate were 2 or 3-fold higher than the B-factors for the CDP portion (Table 1). This indicates the glycerate moieties in these molecules were not well ordered and more dynamic. Overall, this suggests that the structure of CpgD has evolved to preferentially interact with 2-phospho-D-glycerate as a substrate, which is reflected biochemically by the lower K_M,app_^(16)^.

To rationalize the structural basis for the enantiomeric preference, we compared how CpgD interacts with the glycerate moieties in the CDP-2D-glycerate and CDP-2L-glycerate structures (Fig. 5D). The carboxylic acid and hydroxyl groups occupied nearly identical positions in both enantiomers, suggesting that the preference arises from more subtle steric effects. Indeed, inspection of the 2-carbon position revealed a key difference. In the 2D enantiomer, the 2-carbon and its attached hydrogen both point downward, away from the sidechains of Met147 and Lys161, which are positioned above the glycerate moiety. In contrast, in the 2L enantiomer, these atoms both point upward toward Met147 and Lys161. This creates favorable van der Waals interactions in the 2D configuration, with distances of 4.4 and 4.8 Å between the glycerate 2-carbon atom and the carbon atoms of Met147 and Lys161, respectively (Fig. 5D). However, in the 2L enantiomer, the shorter distances of 2.5 and 3.5 Å between the glycerate hydrogen atom and the carbon atoms of Met147 and Lys161 would create steric repulsion (Fig. 5D). Given these differences result directly from the stereochemistry at the 2-carbon position, we concluded that favorable van der Waals interactions in the 2D configuration, together with steric clashes in the 2L enantiomer, would lead to higher affinity of CpgD for 2-phospho-D-glycerate over 2-phospho-L-glycerate and explain its enantiomeric preference.

## DISCUSSION

Here we present a series of high-resolution structures of CpgD in different catalytic states that allowed us to propose a catalytic mechanism for the enzyme and understand the structural basis for nucleotide specificity and phosphoglycerate preference. Favorable hydrogen bonding interactions results in specificity for CTP, while more subtle van der Waals interactions explain the enantiomeric preference for the 2-phospho-D-glycerate substrate. We also captured the conformational changes that occur upon CTP binding, and the lack of any major conformational changes in the product bound state. Overall, this provides a structural basis for the observed *in vitro* substrate specificity and preferences for CpgD.

Despite these advances, we still lack information about the exact chemical structure of the final CPG2 lipid product, for which CpgD and its enzymatic product CDP-2D-glycerate would contribute to. While CpgD displays a preference for the 2-phospho-D-glycerate substrate both biochemically and structurally, CpgD can still utilize 2-phospho-L-glycerate and 3-phospho-D-glycerate as substrates^(16)^. Since the concentrations of these molecules in the cell are unknown, there remains some uncertainty as to which CpgD catalyzed product or products are utilized in CPG2 synthesis. Furthermore, it is unclear if the same enantiomeric preference is shared by the next enzyme in the pathway, CpgA, which presumably uses the product of CpgD catalysis, CDP-glycerate, as a substrate. If CpgA also prefers CDP-2D-glycerate as a substrate over the CDP-2L-glycerate enantiomer, this would suggest that the enzymes involved in CPG2 synthesis have evolved to utilize and incorporate 2-phospho-D-glycerate into CPG2. However, future studies are necessary to resolve this outstanding question.

It is noteworthy that CpgD, which catalyzes the synthesis of CDP-glycerate, shares structural homology with the choline-phosphate cytidylyltransferase that catalyzes CDP-choline synthesis, but no structural homology with the CTP:glycerol-3-phosphate cytidylyltransferases from gram-negative bacteria that catalyze the synthesis of the more closely related molecule CDP-glycerol^(21-23)^. The use of a different protein fold to catalyze similar reactions is in line with the divergent evolution of the bacterial and eukaryotic ceramide synthesis pathways. This not only made it difficult to identify the genes involved in the synthesis of ceramide in bacteria but suggests that *C. crescentus* and other gram-negative bacteria have evolved a distinct set of enzymes, in comparison to the related lipid synthesis pathways, for both the synthesis of the ceramide backbone and the headgroup of CPG2.

Lastly, the role(s) of the additional magnesium ions observed in the product bound structure of CpgD remain unresolved, given they are not observed in the CTP bound structure, and are not present in other structurally characterized nucleotidyltransferases^(18, 19, 24)^. While we hypothesize the third magnesium is likely involved in catalysis by aiding the nucleophilic phosphate attack on the α-phosphate of CTP, we suspect that the fourth observed magnesium ion, which is too far away to directly participate in catalysis, is not an artifact of crystallization. One possible scenario is that this fourth magnesium ion is somehow involved in the selective cooperativity, previously observed when the preferred enantiomer 2-phospho-D-glycerate is used as a substrate. We speculate that this cooperativity may derive from a second 2-phospho-D-glycerate molecule that involves the fourth magnesium binding site. This hypothesis remains speculative given that a second 2-phospho-D-glycerate binding site was not observed in our product bound structure, where the reaction had likely gone to completion. However, time resolved crystallographic studies, which can capture intermediate states of catalysis, may aid in identifying if additional, allosteric 2-phospho-D-glycerate binding sites exist that depend on the fourth magnesium binding site and that can explain the observed cooperativity.

## EXPERIMENTAL PROCEDURES

### Protein expression and purification

CpgD (gene name: *ccna_01210*) was previously cloned into the pET28a expression vector with an N-terminal His-tag^(16)^. For overexpression, the plasmid was transformed into *Escherichia coli* BL21 (DE3) RIPL cells (Agilent Technologies). Cells were grown in 1-liter Terrific Broth (TB) media supplemented with kanamycin (100 μg/mL) at 37 ºC to an optical density at 600 nm of ∼2.5, at which point the incubation temperature was reduced to 15 ºC. After cooling for ∼ 2hrs, IPTG was added to a final concentration of 0.36 mM, and the culture was incubated at 15 ºC overnight for 16-20 h. Cells were harvested by centrifugation, and pellets were stored at -80 ºC.

A frozen cell pellet was thawed and resuspended in lysis buffer adjusted to pH 7.5 (20 mM Tris-HCl pH 7.5, 150 mM NaCl, 10% glycerol, 30 mM imidazole, 10 mM β-mercaptoethanol with the addition of 1 mM protease inhibitors (Aprotinin, Leupeptin, and Pepstatin A). The cells were lysed via sonication at 85% amplitude through repeated cycles of 2 s on/off for 10 mins. The soluble fraction was isolated and insoluble cell debris was removed by centrifugation at 50,000 xg for 30 mins at 4 ºC. The supernatant was loaded into a nickel-nitrilotriacetic acid (Ni-NTA) column after equilibrating with wash buffer (20 mM Tris-HCl pH 7.5, 300 mM NaCl, 30 mM imidazole, and 5 mM β-mercaptoethanol) and eluted in 1 mL fractions using elution buffer (20 mM Tris-HCl pH 7.5, 300 mM NaCl, 300 mM imidazole, and 5 mM β-mercaptoethanol). The protein fractions were directly applied to a size exclusion Superdex 200 pg Hiload 16/600 column (GE Healthcare) with size exclusion buffer (20 mM Tris pH 7.5, 150 mM NaCl, and 10 mM βME). The pooled purified protein was concentrated via centrifugation to 8 mg/mL, aliquoted, flash frozen, and stored at -80°C. Protein concentration was determined by measuring the UV absorption at 280 nm.

### Crystallization and Data Collection

Preliminary crystal hits of apo CpgD were obtained after screening in four 96-well plates of unique crystallization conditions using JSCG Core I-IV suites. The refined growth condition contained 0.1 M sodium citrate, pH 4, and 15% PEG 3350. Crystals were grown in a hanging drop at room temperature with equal volumes of protein solution and buffer solution, which generated large three-dimensional cube-shaped crystals. These crystals were transferred to a cryoprotectant solution containing 40% PEG 3350, 25% glycerol, and 0.1 M sodium citrate, pH 4 prior to freezing and data collection.

Co-crystals of CpgD and CTP were grown by adding 30 mM CTP and 100 mM MgCl_2_ to the protein and the mixture was incubated for 1 hour at 4 °C prior to setting up hanging drops. The refined growth buffer for co-crystals grown with CTP and MgCl_2_ contained 0.1 M Tris pH 7, 20% PEG 400, and 5% glycerol. The cryoprotectant solution for these co-crystals were prepared with 40% PEG 400, 25% glycerol, and 0.1 M Tris, pH 7 and diluted by the addition of CTP and MgCl_2_. The final cryoprotectant solution contained 30 mM CTP, 100 mM MgCl_2_, 0.065 M Tris pH 7, 25% PEG 400, and 15% glycerol.

Co-crystals of CpgD, CTP, MgCl_2_, and 2-phospho-L-glycerate (2-phospho-L-glycerate, Sigma Aldrich catalog# 19710) were grown by adding 30 mM CTP, 30 mM 2-phospho-L-glycerate, and 100 mM MgCl_2_, to the protein and incubated for 1 hour at 4°C. The modified cryoprotectant solution for these co-crystals contained 29.5 mM CTP, 29.7 mM 2-phospho-L-glycerate 95.7 mM MgCl_2_, 22.8% PEG 400, and 14.2% glycerol. Likewise, co-crystals of CpgD with CTP and 2-phospho-D-glycerate (2-phospho-D-glycerate, Sigma Aldrich, catalog# 73885) or 3-phospho-D-glycerate (3-PG, Santa Cruz, catalog# sc-214793) were grown by adding 30 mM CTP, 30 mM 2-phospho-D-glycerate or 3-PG, and 100 mM MgCl_2_, to the protein prior to incubation an hour at 4 °C. The final modified cryoprotectants contained 0.1 M Tris (pH 7 or 7.5), 29.6 mM CTP, 30 mM 2-phospho-D-glycerate or 3-phospho-D-glycerate, 99.4 mM MgCl_2_, 20% PEG 400, and 15% glycerol. Diffraction data were collected at Brookhaven National Lab (BNL) NSLS-II FMX (17-ID-1)^(25)^ or AMX (17-ID-2)^(26)^ beamlines using standby operation^(27)^. Data processing was carried out using the autoPROC pipeline^(28-32)^.

### Structure determination and refinement

Phases for the structure of apo CpgD (CCNA_01210) were determined by molecular replacement (MR) carried out in PHENIX using Phaser. An Alphafold2-predicted structure of CpgD was used as search model, after replacing the B-factor field in the model using Phenix. Phases for CpgD co-crystallized with CTP were determined by MR using apo CpgD structure as a search model. The co-crystal structures with CTP and 2-phospho-D-glycerate were determined by MR using the CTP-bound CpgD structure as a search model. The co-crystal structures with CTP and 2-phospho-L-glycerate and 3-phosphoglycerate were determined by MR using the CDP-2D-glycerate structure as a search model. Ligand restraints for CDP-2D-glycerate, CDP-2L-glycerate, and CDP-3-glycerate were generated using Phenix Elbow^(33)^. All structures were refined using a combination of manual model building in Coot with multiple rounds of refinement in Phenix. Diffraction data collection and refinement statistics are in Table 1. All structural figures were generated using PyMOL 3.1.6.1.

## ACKNOWLEDGEMENTS

The authors thank Drs. Shujuan Gao and Franceine Welcome for their advice and assistance with initial crystallization screens. This research was supported by the NSF grants 2515532 (EAK) and 2515533 (MVA), and in part by the NIH grant T32GM136572 (RR) and an Alfred P. Sloan Fellowship (MVA). This research used resources AMX (17-ID-1) and FMX (17-ID-2) of the National Synchrotron Light Source II, a U.S. Department of Energy (DOE) Office of Science User Facility operated for the DOE Office of Science by Brookhaven National Laboratory under Contract No. DE-SC0012704. The Center for BioMolecular Structure (CBMS) is primarily supported by the National Institutes of Health, National Institute of General Medical Sciences (NIGMS) through a Center Core P30 Grant (P30GM133893), and by the DOE Office of Biological and Environmental Research (KP1605010).

